# Adaptability and yield stability analysis of food barley (*Hordeum vulgare* L.) in South Gondar zone, Northwest Ethiopia

**DOI:** 10.1101/2024.12.31.630857

**Authors:** Alamir Ayenew Worku, Dejen Bekis Fentie, Solomon Sharie Shferaw

## Abstract

In food barley, genotype-environment interaction is a frequent occurrence. By identifying stable genotypes across settings, the impact of G x E can be mitigated. Ten food barley genotypes were investigated for yield stability and adaptation in five different environments. The analyses included stability, genotype, and genotype-environment interaction, as well as additive main effect and multiplicative interaction. However, genotype-environment interaction captured 92.98% of the GGE variance. AMMI analysis only explained 27.87% of G and 20.12% of G x E. Five environments were identified using GGE analysis. Of these, Estie2022 (3591.3 kg ha^−1^) and Debretabor2022 (3494.47 kg ha^−1^) had the highest grain yields, indicating that they are suitable for the production of food barley, while Lay-Gaynt2023 (2399.82 kg ha^−1^) had the lowest grain yields, indicating that they are not a suitable environment. The conditions with the best yields were Estie2022, and the lowest-yielding environments were Lay-Gaynt 2023. The two genotypes with the highest yields were HB-1966 (3592.97 kg ha-1) and HB-1965 (3555.17 kg ha-1). Based on yield stability, the genotypes Biftu, followed by Adoshe and Abdene, were the most stable with the highest mean grain yield performance. The genotypes with the highest-yielding performance but relatively the lowest stability were HB-1966 and HB-1965. The genotype HB-1965 was recommended at the Estie2022 and Debretabor2023 testing environments. The ideal environment was represented by Estie2023, and the winning genotype was HB-1966. This suggests that the two winning genotypes were adapted to the specific environments tested in the current study. Whereas Guta and Harbu were adapted to a more diverse environment than the test environments of the South Gondar Zone, Northwest Ethiopia.

## Introduction

Barley (Hordeum vulgare L.) belongs to the genus Hordeum in the Triticaceae of the Gramineae family. It is self-pollinated diploid 2n=14. In Ethiopia, barley is the fifth most important cereal crop after tef, maize, sorghum, and wheat in both total area coverage and annual production (Bakala et al., 2022). The total area covered by barley in Ethiopia is about 959,273.40. ha, with a total production of 2,024,921.70 tons, and its productivity is 2.11 tons/ha (Belete, 2023). The total area covered by barley in the Amhara region is about 339,689. hectares, with a total production of 633,115.4 tons, and its productivity (1.86 tons/ha) is below the national average of 2.11 tons/ha (Birhanu et al., 2020).

Barley is a main food crop in the highland areas where other cereals cannot grow, as well as animal feed and forage around the world. Barley grain contains 3 to 7% β-glucan, which is an important dietary fiber that has significant blood cholesterol-dropping effects. It also plays an important role in ensuring food security (Hailu et al., 2015).

Ethiopia’s food production faces several challenges, including a lack of widely adapted, high-yielding, disease- and insect-resistant varieties; seasonal and regional weather variations; declining soil fertility; and the growing effects of climate change. However, evaluating food barley types in the research locations has not been extensively studied (Shibeshi and Mekiso, 2022).

In food barley, stability analysis assesses a cultivar’s ability to perform consistently in various environmental conditions. It assists breeders in choosing cultivars that consistently produce well under various circumstances. Stability research finds cultivars that show broad adaptability by taking into consideration different locales and seasons, guaranteeing constant production of food barley (Sayed et al., 2018).

The stability of agricultural cultivars under varied environmental conditions is assessed using a set of analytical tools known as stability statistics. These statistics include the mean square of deviations from the regression (S^2^di), the coefficient of determination (R^2^), the ecovalence (Wi) statistic, and the genotype main effect plus genotype-by-environment interaction (GGE) biplot analysis. Because they provide information on the genotype-by-environment interaction, they can help breeders select stable cultivars with reliable performance (Sayed et al., 2018). For food barley varieties that will improve both short-term high grain output and long-term food security, high-yielding potential, enhanced stability, and adaptability are essential. It is important to focus on finding more stable varieties, particularly in Ethiopia, where it is costly to produce highly adapted cultivars. The current study’s objectives were to identify high-yielding variations that were extensively and particularly adapted and to evaluate the long-term dependability of food barley genotypes in the South Gondar zone of Northwest Ethiopia.

## Materials and Methods

### Planting materials, experimental design, and test locations

Ten improved food barley varieties were brought from the Sinanna Agricultural Research Center and evaluated at three locations with five distinct conditions during the cropping season of 2022-2023 (Table 2). The soil type, altitude, temperature, and annual rainfall obtained in every single location differ (Table 1). At every location, a randomized complete block design was used with three replications. The plot area was 3 m^2^ (1.2 m × 2.5 m), with 1.5 m, 0.5 m, and 0.2 m distances between the plot, row, and replications, respectively. 125 kg/ha of seed was sown using the drilling method, and fertilizer was applied in the amounts of 41 N and 46 P_2_O_5_ kg ha-1 of urea and NPS.

**Table 1.** Description of experimental locations.

| Location | Elevation(m) | latitude | longitude | Annual<br>rainfalls<br>(mm) | Temperatur<br>e (° C) | Soil type |
| --- | --- | --- | --- | --- | --- | --- |
| Debre tabor | 2630 | 11.87 °N | 38.03 ° E | 1553.03 | NA | Luvisol |
| Estie | 2268 | 12.95 °N | 38.72 ° E | 1220 | 20.5 | Red |
| Lay -Gaynt | 2700 | 12.00 °N | 38.19 ° E | 997 | 12.84 | Red |

NA=Not available

**Table 2.** Genotypes used in the study.

| S. N | Genotype code | Genotype Name | Year of release | Research center |
| --- | --- | --- | --- | --- |
| 1 | G1 | Biftu | 2005 | SARC |
| 2 | G2 | Robera | 2016 | SARC |
| 3 | G3 | HB-1965 | 2017 | HARC |
| 4 | G4 | Abdane | 2011 | SARC |
| 5 | G5 | Dinsho | 2004 | SARC |
| 6 | G6 | Adoshe | 2018 | SARC |
| 7 | G7 | Defo | 2005 | SARC |
| 8 | G8 | HB-1966 | 2017 | HARC |
| 9 | G9 | Guta | 2007 | SARC |
| 10 | G10 | Harbu | 2004 | SARC |

**S. N= Serial Number, SARC=Sinana Agricultural Research Center, HARC=Holeta Agricultural Research Center**

### Data collection

Phenological and morphological characteristics were determined according to barley descriptors (IPGRI, 1994) based on plant-based and plot-based traits. For plant-based traits plant height, number of grains per spike, and spike length were considered. Ten randomly selected plants from the central part of the row were tagged before data collection and measured timely according to the traits nature to use. For plot-based traits, thousand-seed weight and grain yield per hectare were taken from the whole row for analysis. In addition, days to heading and days to maturity were counted from emergence to 50% heading and 90% physiological maturity.

### Statistical analysis

Analysis of variance, for the combined data across locations and over years, was carried out using R4.4.0 software. Mean separation was carried out using the least significant difference (LSD) at a 5% level of significance. The G x E interaction was further partitioned using additive main effects and multiplicative interaction statistical model by R software. The AMMI analysis of variance summarizes most of the magnitude of genotype x environment interaction into one or more interactions principal component analysis (Zobel *et al*., 1988; Guach, 1988). The larger the IPCA scores, the more specifically adapted a genotype is to a certain environment whereas the smaller the IPCA scores, the more stable the genotype in studied environments. Eberhart and Russell (1966) computed the regression coefficient (bi), deviation from regression (Sdi^2^), and coefficient of determination (R^2^). It was calculated by regressing the mean grain yield of individual genotypes on the environmental index. Shukula stability variance (σ^2^) and Ecovalence (Wi) computed by Wricke (1962), where the low values are considered stable. Cultivar superiority was computed by (Lin and Binns, 1988). Variability of genotype was also measured by coefficient of variation (CV) computed by Francis and Kannenberg (1978) and genotypic variance across environmental (Si^2^).

## Results and discussion

### Analysis of variance

The analysis of variance showed a highly significant (P < 0.01) difference between genotypes for the traits considered in the experiment (Table 3). Locations have a highly significant difference (p<0.01) for days to heading, days to maturity, plant height, biomass yield, and harvest index. The year also showed a highly significant difference (p<0.01) for days to heading, days to maturity, plant height, spike length, biomass yield, grain yield, and harvest index. The G x L was highly significant (P < 0.01) for days to heading, days to maturity, plant height, and number of grains per spike. While G x Y was highly significant (P < 0.01) for days to maturity, plant height, number of grains per spike, biomass yield, grain yield, and harvest index. G x L x Y effect was highly significant (P < 0.01) for days to maturity, number of grains per spike, and significant difference (p<0.05) for grain yield. According to Birhanu et al.,2020 found significant differences in Genotype, Location, and genotype by location interaction effects that there was a highly significant difference at (p≤0.01) among varieties for days to maturity, number of grains per spike, and grain yield. Birhanu et al., (2020) obtained significant differences in location (P < 0.01) for days to heading, days to maturity, and plant height. Buli, (2023) reported highly significant differences at (p<0.01) for G, L, G x L on plant height and spike length.

**Table 3.** Analysis of variance of 10 food barley genotypes tested at three locations for two years.

| Genotypes | DH | DM | PH | SL | NGS | BY | GY | TSW | HI |
| --- | --- | --- | --- | --- | --- | --- | --- | --- | --- |
| Genotype (G) | ** | ** | ** | ** | ** | ** | *** | ** | ** |
| Location (L) | ** | ** | ** | * | ns | ** | ns | ns | ** |
| Year (Y) | ** | ** | ** | ** | ns | ** | ** | ns | ** |
| G x L | ** | ** | * | ns | ** | ns | ns | * | ns |
| G x Y | ns | ** | ** | ns | ** | ** | ** | ns | ** |
| G x L x Y | ns | ** | * | ns | ** | ns | * | ns | ns |

\*\* and \* = Significant at 0.01 and 0.05 probability levels, respectively and <sup>ns</sup>=non-significant difference, **DH**=days to heading, **DM**=days to maturity, **PH**=plant height, **SL**= spike length, **NGS**=number of grains per spike, **BY**=biomass yield, **GY**= Grain Yield, **TSW**= Thousand seed weight, **HI**= harvest index

### Performance of the genotypes

The mean values of the food barley genotypes for the traits considered are depicted in Table 4. Genotypes G8 and G3 were the first and second highest-yielding genotypes, with yields of 3592.97 and 3555.17 kg ha1, respectively, while Guta (G9; 2426.23 kg ha1) was the least. According to thousand-grain weight, HB-1966 (40.01 g) were comparatively larger seeded genotypes, whereas Defo (G7; 33.21 g) and G5 (37.13 g) were small-seeded types. Likewise, Biftu (102.95 cm), followed by G4 (100.69 cm), and G10 (100.19 cm), were the tallest, whereas G6 (83.67 cm) and G3 (93.03 cm) were shorter genotypes. HB-1966 (G3), Adoshe (G6), and HB-1965 (G8) took 75.47, 76.00, and 74.8 days to head, while G9, G7, G5, and G10 took 66.33, 67.87, 67.93, and 68.13 days, respectively. In most cases, days to maturity were also similar to days to heading (Worede *et al*., 2020). HB-1966 (G3), Adoshe (G6), and HB-1965 (G3) took 122.67, 116.67, and 115.6 days, respectively, while Guta (G9), Defo (G7), and Habiru (G10) took 106.73, 107.2, and 108.47 days, respectively.

**Table 4.**
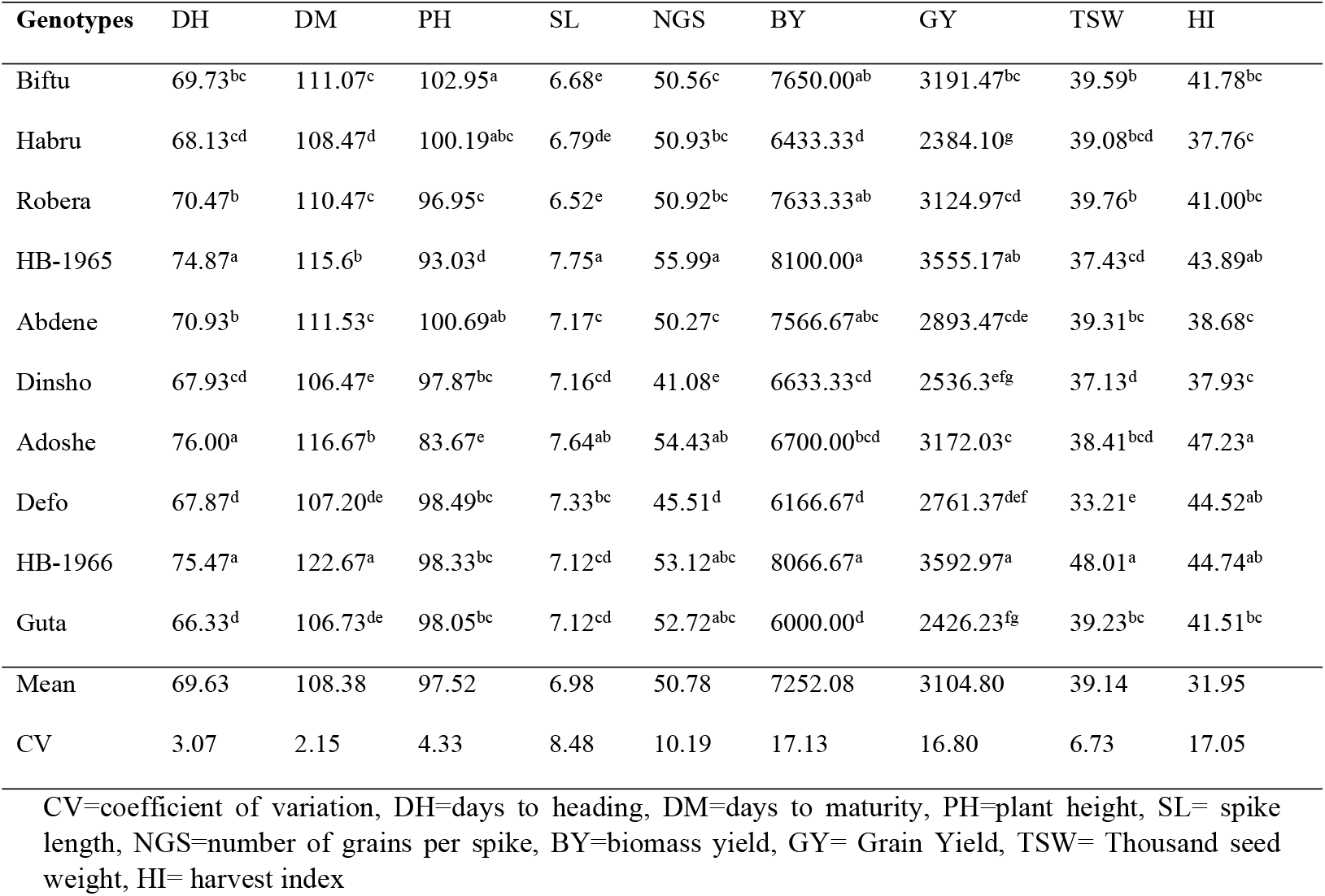
Mean grain yield and other agronomic traits of food barley genotypes tested at three locations for two years.

Estie 2022 (3591.3 kg ha^−1^), Debretabor 2022 (3494.47 kg ha^−1^), and Lay-Gaynt 2023 (2399.82 kg ha^−1^) had the highest and lowest food barley grain yields, respectively, when the grand mean values of the five environments were calculated. One could consider Lay-Gaynt 2023 to be the lowest-yielding environment and Estie 2022 to be the highest. The genotypes with the best yields were G3 (3555.17 kg ha^−1^) and HB-1966 (3592.97 kg ha^−1^). Conversely, the genotypes with the lowest yields were Guta (2426.23) and G10 (2384.1 kg ha^−1^) (Table 5).

**Table 5.** Mean grain yield (kg/ha) of 10 food barley genotypes across five environments (location and year combinations)

| **Genotype** | **E1** | **E2** | **E3** | **E4** | **E5** | **Mean** |
| --- | --- | --- | --- | --- | --- | --- |
| **Biftu** | 3680.83 | 3659.67 | 3206.00 | 2845.00 | 2565.83 | **3191.47** |
| **Habru** | 3352.83 | 3109.33 | 1786.00 | 1500.67 | 2171.67 | **2384.10** |
| **Robera** | 3707.50 | 4139.17 | 3418.50 | 2173.67 | 2186.00 | **3124.97** |
| **HB-1965** | 3314.67 | 4420.33 | 4151.00 | 3190.67 | 2699.17 | **3555.17** |
| **Abdene** | 3288.17 | 3054.00 | 2889.33 | 2736.50 | 2499.33 | **2893.47** |
| **Dinsho** | 3504.50 | 2968.33 | 2005.33 | 1659.50 | 2543.83 | **2536.30** |
| **Adoshe** | 3179.67 | 4111.17 | 3026.17 | 3285.00 | 2258.17 | **3172.04** |
| **Defo** | 3774.33 | 3390.50 | 2017.00 | 2190.00 | 2435.00 | **2761.37** |
| **HB-1966** | 3777.33 | 4161.67 | 3321.00 | 4474.67 | 2230.17 | **3592.97** |
| **Guta** | 3364.83 | 2898.83 | 1754.50 | 1704.00 | 2409.00 | **2426.23** |
| **Mean** | **3494.47** | **3591.30** | **2757.48** | **2575.97** | **2399.82** | **2963.809** |
| **CV** | 12.35 | 11.39 | 18.38 | 18.41 | 29.69 |  |
| **LSD** | **ns** | 701.87 | 869.28 | 813.30 | **ns** |  |

CV=coefficient of variation, LSD=Least Significant Difference, E1= Debretabor2022, E2= Estie2022, E3= Debretabore2023, E4= Estie2023, E5= Lay- Gaynt2023.

### AMMI analysis

AMMI analysis of variance revealed a significant difference (P < 0.01) between the genotypes (G) and the interaction (G x E). Additionally, 27.87 and 20.12% of the treatment sum of squares (SS) were accounted for by the G and G x E components, respectively. Only the G and G x E factors are relevant because they influence the genotype ranking, even though the genotypes explained the majority of the variance in the current study (Gauch & Zobel, 1997). Additionally, one IPCA (Interaction Principal Component Axes) was generated from the G x E, and it explained 14.05% of the G x E and was significant (P < 0.01) (Table 7).

The AMMI biplot (Figure 1) showed that G8 and G3 were the genotypes with higher yields, whereas G1 and G6 had above-average grain yields. The grain yield of genotype G2 was equal to the grand mean value, whereas the grain yields of genotypes G10, G9, G5, G7, and G4 were below average. But G10 and G9 were the lowest yielders. Furthermore, in G8, G3, G9, G5, and G10, a substantial genotype by environment interaction was seen. On the other hand, G1’s G x E was moderate, while G2, G4, and G4’s G x E were lower.

**Figure 1.**
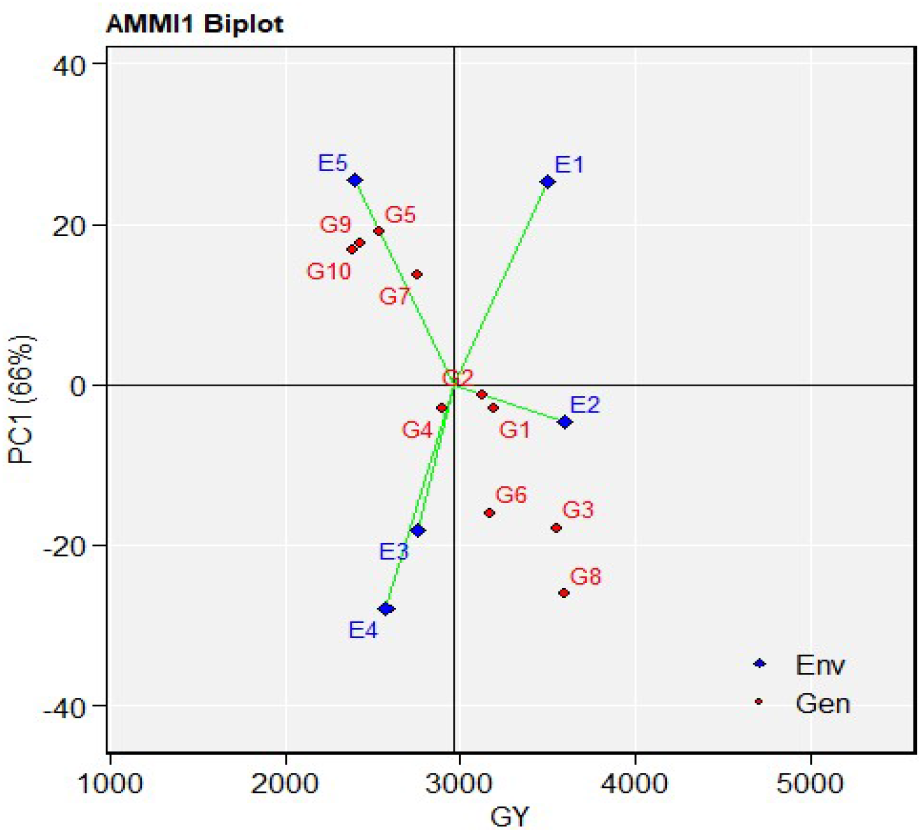
AMMI 1 biplot of main effects of food barley genotypes and environments, and IPCA1 using symmetrical scaling, E1= Debretabor2022, E2= Estie2022, E3= Debretabore2023, E4= Estie2023, E5= Lay-Gaynt2023.

### Genotype and genotype by environment interaction (GGE) biplot analyses

Sidelines connect the environmental ratings to the origin (Figure 2). Weak contact forces have been exerted in positions with short bars. Long bar users engage in intense interaction. Therefore, environments Debretabor2022 and E5 exerted weak contact forces, while Estie2022, E3, and E4 exerted significant ones. Due to their distance from the origin, genotypes HB-1966, G3, and G2 in this study were more responsive to environmental interaction forces than genotypes G6, G4, Defo, and G9, which were less sensitive. The genotypes G5, G10, and Biftu, on the other hand, were the closest to the origin and had almost no interaction forces.

**Figure 2.**
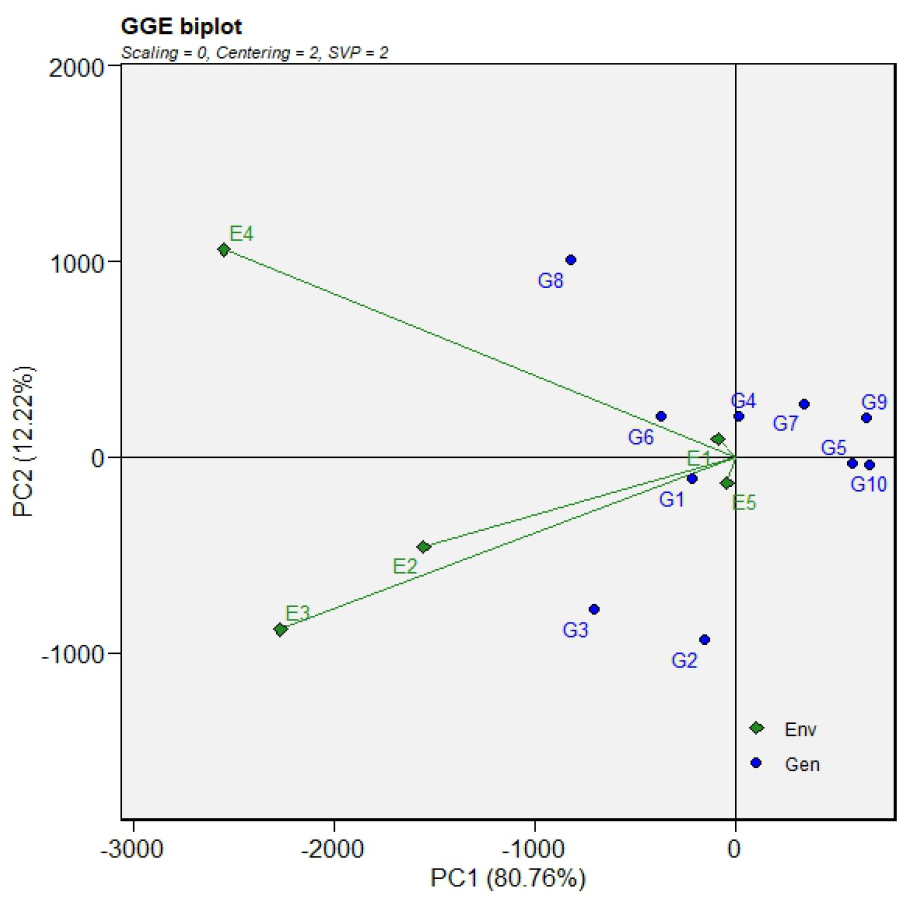
GGE biplot for grain yield (kg ha^−1^) showing the interaction of PC1 against PC2 scores of Ten food barley genotypes (G) and five environments(E), E1= Debretabor2022, E2= Estie2022, E3= Debretabore2023, E4= Estie2023, E5= Lay-Gaynt2023

### Mean grain yield and stability performance of genotypes

The environment coordination (AEC) approach is used to graphically depict the genotypes’ mean grain yield and stability performance (Figure 3). The most stable genotypes producing the highest and most stable yields can be found using this method, which combines genotype stability performance with grain yield. The optimum genotype is the one with the best mean performance and the highest stability across all test settings (Seyoum, 2021).

**Figure 3.**
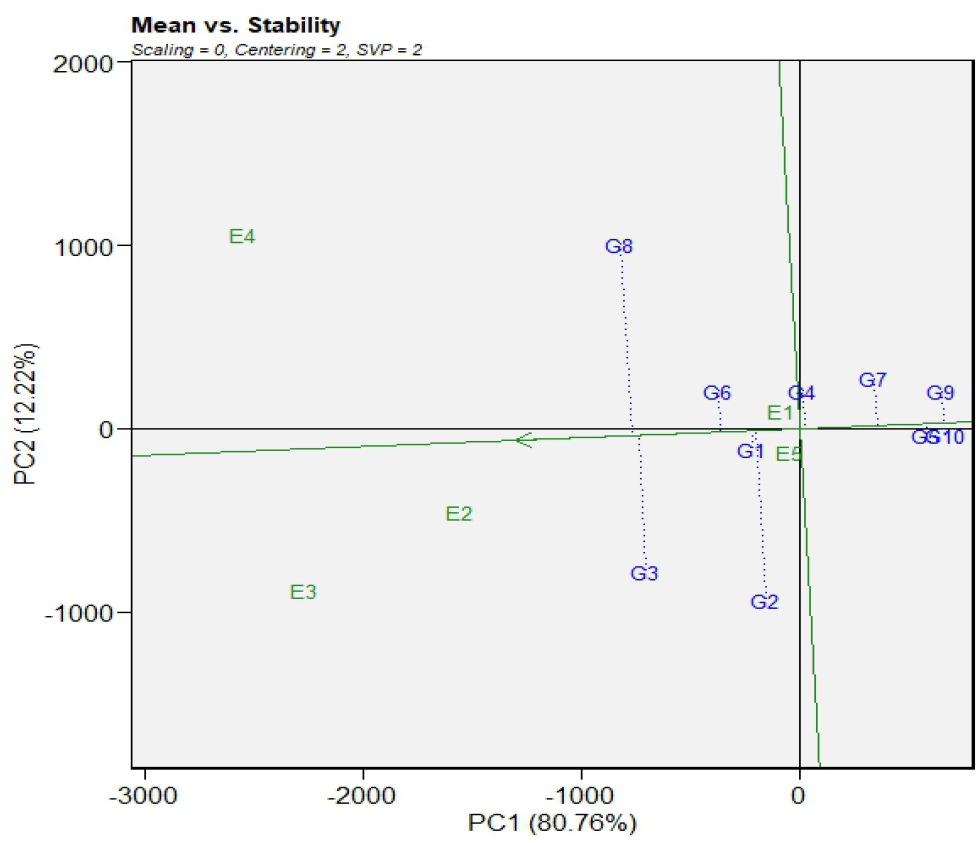
Ranking genotypes based on mean grain yield and stability across environments E1= Debretabor2022, E2= Estie2022, E3= Debretabore2023, E4= Estie2023, E5= Lay-Gaynt2023

A desirable genotype is located closer to an ideal genotype, which is typically at the center of the concentric circles or arrows, according to the AEC view comparison biplot. The high-yielding genotypes with the longest vector length are linked to an optimum genotype. An arrow on the AEC X-axis (PC1), which passes through the biplot origin in the average environmental coordinate (AEC) system, represents the mean performance axis of genotypes. The biplot origin and the ATC Y-axis are both perpendicular to one another. The stability axis (PC2) is indicated by this axis. On the positive side of the AEC abscissa, genotypes located closer to the origin would have a higher mean grain yield. In contrast, genotypes located farther away on the negative side would have a lower mean grain yield. Additionally, a genotype’s projection becomes less stable the longer it is in absolute terms. This line points towards a higher mean yield. across the environment. Hence, in the present study, G8 gave the highest mean yield followed by G3 and G2 (Figure 3). The line that passes through the biplot and is perpendicular to the AEC axis shows the measure of stability. Greater genotype x environment interaction and poor stability are indicated by either direction on this axis away from the biplot origin, or vice versa. In terms of stability, the genotype ranks as G5>G10>G1>G9>G4>G6>G3>G8 (Figure 3).

The estimates of eight stability coefficients for the 10 food barley genotypes computed based on five environments (location and year combination) are displayed in Table 7. Based on cultivar superiority stability statistics, HB-1966 (15.15) and HB-1965 (27.48) were the most stable, as they had comparatively smaller values; G10 (245.27) and G9 (225.07) were the least stable.

A relatively large stability variance value (σ^2^) indicates greater instability of the genotypes. Thus, the genotypes Biftu and Abdene with a low value of stability variance and mean grain yield above the average were stable, and the genotypes HB-1966 (133.39), G2 (36.16), HB-1965 (35.57), G5 (36.65), and Guta (33.8) with a high stability variance were considered unstable (Table 8).

A genotype with Wi = 0 is considered stable, based on Wricke (1962). Accordingly, Abdene (0.03), Biftu (0.68), Defo (5.14), and G6 (6.17) were the most stable. G8 with a value of 37.5, G5 with 11.1, and G2 with a value of 11.0 were the least stable (Table 7).

The coefficient of determination (R^2^) represents agronomic stability (Becker, 1981). A genotype with a high coefficient of determination can be considered stable. The expectedness of genotypes for the grain yield ranged from 0.23 to 0.99, indicating that 23–99% of the mean grain yield variation was explained by genotype responses across different environments. Genotypes with a low coefficient of determination were considered unstable. The genotype HB-1966 (0.23) had low values considered unstable, whereas the genotypes Biftu and Defo (0.99), followed by Abdene (0.98), had high values considered stable (Table 6).

**Table 6.** The various models of stability used to partition the G x E for grain yield in the test food barley varieties and their ranking.

| Genotypes | Mean | bi | s<sup>2</sup>di | R<sup>2</sup> | $\sigma^2$ | wi | pi | s<sup>1</sup> | s<sup>2</sup> |
| --- | --- | --- | --- | --- | --- | --- | --- | --- | --- |
| Biftu | 2705 | 1.79 | -10.91 | 0.99 | -1.52 | 0.68 | 59.44 | 0.33 | 1.00 |
| Habru | 1836 | -1.06 | 4.43 | 0.32 | 22.61 | 7.28 | 245.27 | 1.67 | 8.33 |
| Robera | 2180 | 3.46 | 14.58 | 0.75 | 36.16 | 11.00 | 101.58 | 0.33 | 14.3 |
| HB-1965 | 2945 | 4.07 | -7.66 | 0.97 | 35.57 | 10.80 | 27.48 | 1.00 | 12.3 |
| Abdene | 2618 | 1.09 | -10.86 | 0.98 | -3.96 | 0.03 | 77.55 | 1.67 | 16.3 |
| Dinsho | 2102 | -1.49 | 14.56 | 0.36 | 36.65 | 11.10 | 209.22 | 2.33 | 14.3 |
| Adoshe | 2772 | 2.13 | 17.05 | 0.51 | 18.55 | 6.17 | 47.92 | 1.67 | 13.00 |
| Defo | 2312 | -1.17 | -10.89 | 0.99 | 14.77 | 5.14 | 164.06 | 0.33 | 14.30 |
| HB-1966 | 3352 | 3.00 | 183.31 | 0.23 | 133.39 | 37.50 | 15.15 | 2.00 | 12.00 |
| Guta | 2056 | -1.82 | -1.23 | 0.68 | 33.80 | 10.30 | 225.07 | 2.67 | 16.00 |

$R^2$ =Coefficient of determination, $S^1$ = Nassar and Huens's (1987) Mean of the absolute rank differences of a genotype over the n environments, $S^2$ = Nassar and Huens's (1987) variance among the ranks over the k environments, $W_i$ =Wriecke's ecovalence, $P_i$ =Cultivar superiority, $\sigma^2$ = Shukla's (1972) stability variance, $b_i$ =Eberhart and Russell (1966) stability value of regression coefficient and $S^2d_i$ = Eberhart and Russell (1966) stability deviation value from regression.

**Table 7.** AMMI analysis of variance for grain yield of 10 food barley varieties.

| Source of variation | DF | Sum Square | Mean Square | % of Explained SS |
| --- | --- | --- | --- | --- |
| Environments | 2 | 1919026 | 959513<sup>ns</sup> |  |
| Replication (Environment) | 6 | 8175488 | 1362581.3** | **9.34** |
| Genotypes | 9 | 24387700 | 2709744.4** | **27.87** |
| Genotype x Environments | 18 | 17605696 | 978094.2** | **20.12** |
| IPC1 | 10 | 12290060 | 1229006** | **14.05** |
| IPC2 | 8 | 5315636 | 664454.5<sup>ns</sup> |  |
| Residuals | 54 | 17808715 | 329791.0 |  |
| Total | 107 | 87502320 | 817778.6 |  |

\*\* = Significant at 0.01 probability level and <sup>ns</sup>=non-significant, DF=degree of freedom, SS=sum of square.

According to the Eberhart and Russell model, regression coefficients and deviation from regression ranged from −1.06 to 4.09 and from 4.43 to 17.05, respectively, indicating that genotypes already had different responses to environmental changes. According to this model, a stable genotype should have a high mean yield (b = 1.0 and S2 = 0). Thus, the genotypes G10 (−1.06), Dinsho (−1.49), Defo (−1.17), and Guta (−1.8) and Abdene (1.09) with regression coefficients close to 1 and Guta (−1.23) and Habru deviation from regression close to zero were considered stable (Table 6).

Genotypes with less change in rank are expected to be more stable. The mean absolute rank difference (S^1^) estimates are all possible pair-wise rank differences across locations for each genotype. The S^2^ estimates are simply the variances of ranks for each genotype over environments (Huhn, 1996). For S^1^, entries may be tested for being significantly less or more stable than the average. Usually, S^1^ is the preferred parameter because of its ease of computation and clear and relevant interpretation. Biftu (1.00) and Habru (8.33) had the lowest variance of rank differences, and they were stable genotypes (Table 6).

### Which-won-where, GGE biplot, and environment relationship

The which-won-where view of the GGE biplot explained 92.47% of the total variation, 60.76% of which was explained by PC1 and 21.71% by PC2 (Figure 4). The genotypes in the vertex (five genotypes) formed a five-sided polygon. The lines drawn from the center perpendicular to the sides of the polygon divided the biplot into five sectors, and the five environments fell into three of the five sectors. For environments within a sector, the winning genotype was located at the vertex of the sector. The genotype HB-1965 was recommended at Estie2022 and Debretabor2023 testing environments. The ideal environment was represented by Estie2023, and the winner genotype was HB-1966. This suggests that the two winning genotypes were adapted to the specific environments tested in the current study. Guta and Harbu were adapted to a diverse environment than the test environments (Figure 4).

**Figure 4.**
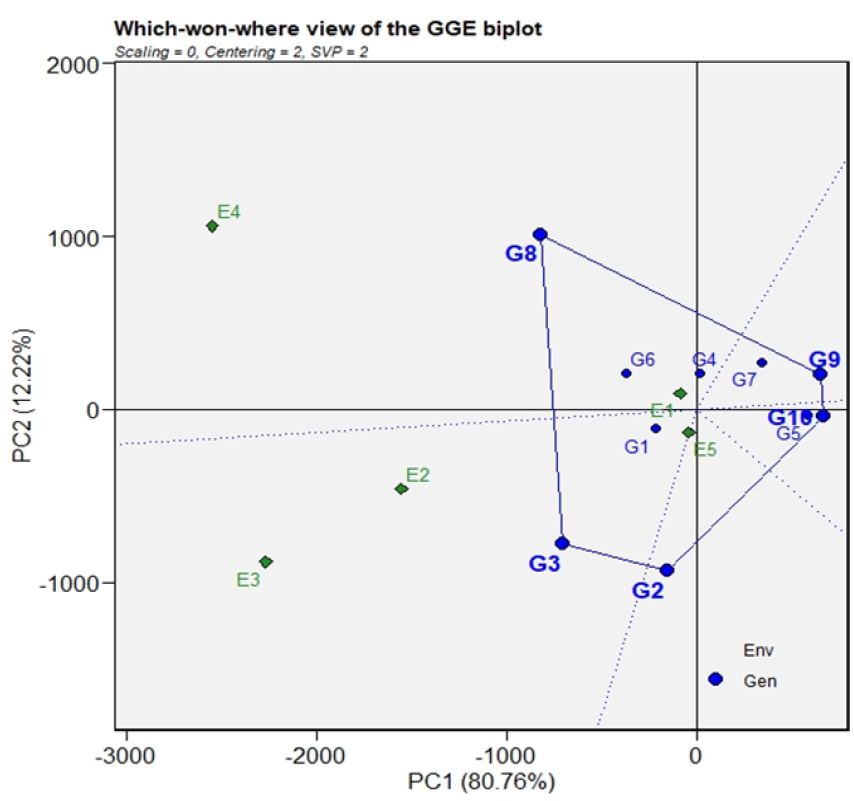
GGE-biplot showing food barley genotypes concerning the environments, E1= Debretabor2022, E2= Estie2022, E3= Debretabore2023, E4= Estie2023, E5= Lay-Gaynt2023.

### Environment ranking for all genotypes

Through the representation of the center of the concentric circles in a polygon perspective, the discrimination and representative of the GGE biplot often demonstrate the ranking of an ideal test environment for a particular genotype or all genotypes (Figure 5). A point on the average environment axis pointing in a positive direction, referred to as “most representative,” with the longest environment vector from the biplot origin is considered the optimal test environment in the GGE biplot (Yan and Tinker 2006). The test environment that is most representational of the target environment and most discriminating (informative) is the optimal one. The ideal test environment is represented by the middle of the concentric circles (Figure 5). Debretabor2022 and Lay-Gaynt2023 performed the worst when it came to choosing genotypes that were adapted to the entire region, whereas Debretabor2023 is the closest to the optimal environment point.

**Figure 5.**
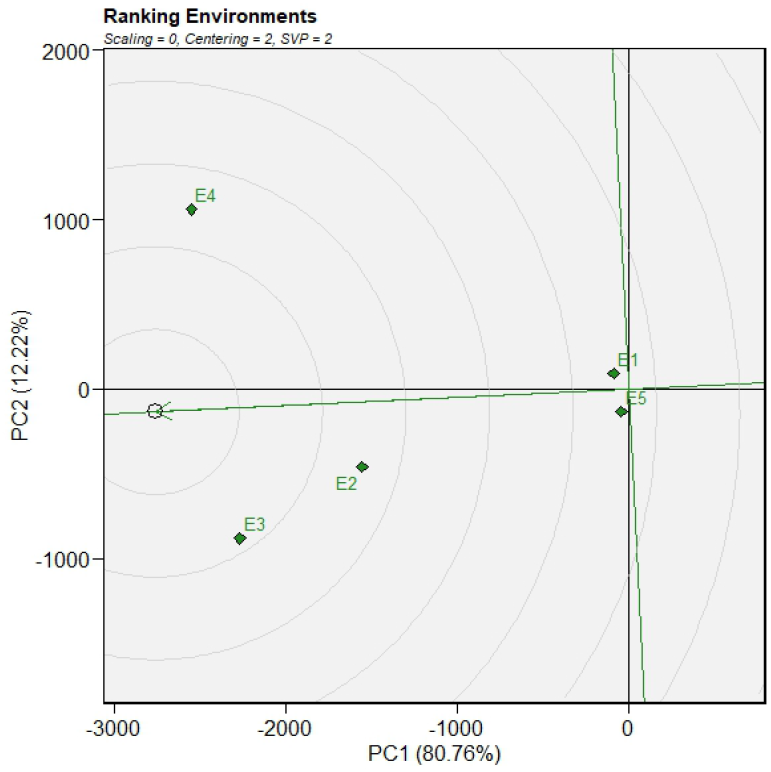
GGE biplot view to rank the five-food barley growing environments using environment-centered scaling. E1= Debretabor2022, E2= Estie2022, E3= Debretabore2023, E4= Estie2023, E5= Lay-Gaynt2023.

### Genotype ranking for all environments

In the GGE biplot analysis, an ideal genotype is a virtual genotype that should have both a high mean yield and high stability. For this purpose, the origin and average point of genotypes are connected and continue to both ends (Maniruzzaman *et al*.,2019). The position of the ideal genotype is at the center of the concentric circles (Figure 6). In the current study, the ranking of genotypes was shown by the comparison with the ideal genotype. The best genotype is a genotype that is closer to the ideal genotype position. Thus, genotypes ‘G1’ and ‘G6’ which were close to the ideal genotype position, were considered ideal genotypes in our study in terms of yield capacity and stability compared to other genotypes.

**Figure 6.**
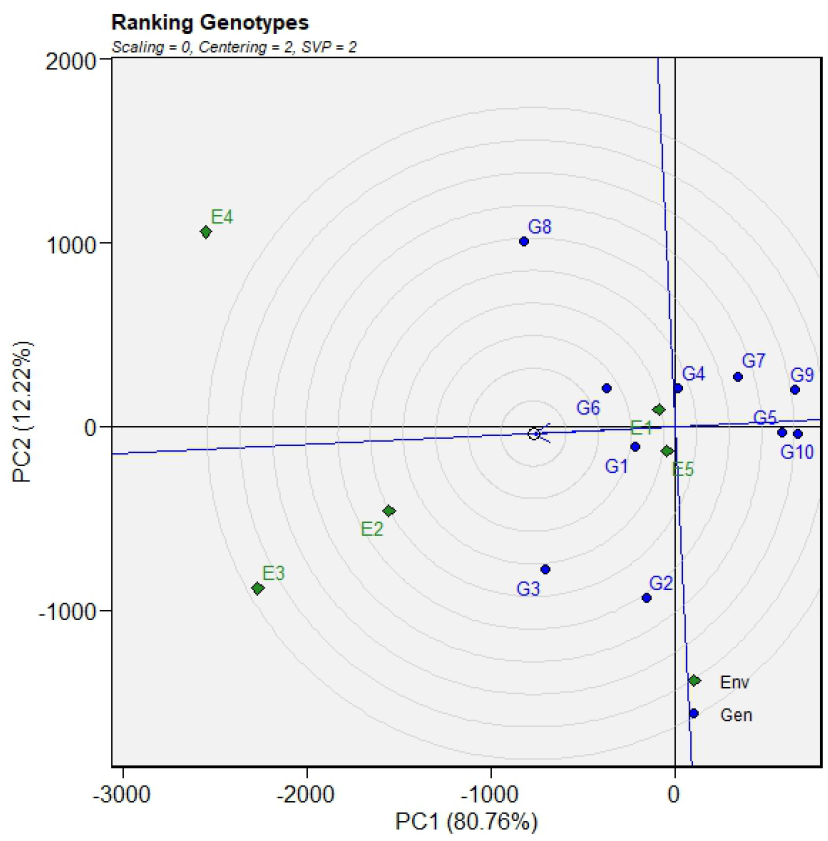
GGE biplot of food barley genotypes on five-environments using genotype-centered scaling. E1= Debretabor2022, E2= Estie2022, E3= Debretabore2023, E4= Estie2023, E5= Lay-Gaynt2023.

## Conclusion and recommendation

food barley, a resilient crop with high nutritional value, plays a vital role in global food security due to its adaptability and disease resistance. Screening and shuttle breeding in contrasting environments, followed by multi-locational evaluations, are key to identifying stable, high-yield genotypes. HB-1966 (3592.97 kg ha^−1^) and HB-1965 (3555.17 kg ha^−1^) were the top-yielding genotypes, while Harbu (2384.1 kg ha^−1^) and Guta (2426.23 kg ha^−1^) had the lowest yields. Biftu, Adoshe, and Abdene showed the highest yield stability, whereas HB-1966 and HB-1965 combined high yields with lower stability. HB-1965 performed best at Estie2022 and Debretabor2023, while HB-1966 was excellent at Estie2023 testing environments. Guta and Harbu demonstrated broader adaptability beyond Northwest Ethiopia.

## Data availability statement

Data will be made available on request.

## Credit authorship contribution statement

**Alamir Ayenew:** Writing review and editing, Writing Original draft, Visualization, Validation, Supervision, Software, Resources, Project administration, Methodology, Investigation, Formal analysis, Data curation, Conceptualization. **Dejen Bekis:** Writing review and editing, Supervision. **Solomon Sharie:** Data curation and visualization.

## Acknowledgments

This research was funded by the Ethiopian Institute of Agricultural Research (EIAR). We acknowledge the Fogera National Rice Research and Training Center and the Debre-Tabor Agricultural Research Sub-center for providing essential experimental equipment and facilitating budget support.

